# Clinical and molecular features of acquired resistance to immunotherapy in non-small cell lung cancer

**DOI:** 10.1101/2021.07.21.452854

**Authors:** Danish Memon, Hira Rizvi, George Fromm, Jayon Lihm, Adam J. Schoenfeld, Jennifer L. Sauter, Jia Luo, Andrew Chow, Umesh K. Bhanot, Caroline McCarthy, Darwin Ye, Chad M. Vanderbilt, Cailian Liu, Mohsen Abu-Akeel, Andrew J. Plodkowski, Nicholas McGranahan, Marta Łuksza, Benjamin D. Greenbaum, Taha Merghoub, Andy J. Minn, Pedro Beltrao, Taylor H. Schreiber, Martin L. Miller, Matthew D. Hellmann

**Author notes:** Contributed equally (Martin L. Miller, Matthew D. Hellmann). Correspondence: Martin L Miller, Matthew D. Hellmann.

## Abstract

Although cancer immunotherapy with PD-(L)1 blockade is now routine treatment for patients with lung cancer, remarkably little is known about acquired resistance. We examined 1,201 patients with NSCLC treated with PD-(L)1 blockade to clinically characterize acquired resistance, finding it to be common (occurring in more than 60% of initial responders), with persistent but diminishing risk over time, and with distinct metastatic and survival patterns compared to primary resistance. To examine the molecular phenotype and potential mechanisms of acquired resistance, we performed whole transcriptome and exome tumor profiling in a subset of NSCLC patients (n=29) with acquired resistance. Systematic immunogenomic analysis revealed that tumors with acquired resistance generally had enriched signals of inflammation (including IFNγ signaling and inferred CD8+ T cells) and could be separated into IFNγ upregulated and stable subsets. IFNγ upregulated tumors had putative routes of resistance with signatures of dysfunctional interferon signaling and mutations in antigen presentation genes. Transcriptomic profiling of cancer cells from a murine model of acquired resistance to PD-(L)1 blockade also showed evidence of dysfunctional interferon signaling and acquired insensitivity to *in vitro* interferon gamma treatment. In summary, we characterized clinical and molecular features of acquired resistance to PD-(L)1 blockade in NSCLC and found evidence of ongoing but dysfunctional IFN response. The persistently inflamed, rather than excluded or deserted, tumor microenvironment of acquired resistance informs therapeutic strategies to effectively reprogram and reverse acquired resistance.

## Introduction

PD-(L)1 blockade can generate profound, durable responses in patients with lung cancer and has been rapidly incorporated into the treatment paradigm for nearly all patients with advanced non-small cell lung cancer (NSCLC) (Gandhi et al., 2018; Reck et al., 2016). Unfortunately, even among those patients who initially respond to PD-(L)1 blockade, over half will eventually develop progression – termed acquired resistance (Schoenfeld and Hellmann, 2020). Alongside primary resistance (refractory to initial treatment), acquired resistance represents a significant and possibly underappreciated clinical challenge (Garon et al., 2019; Gettinger et al., 2018a, 2018b; Herbst et al., 2020). Remarkably little is known about the molecular mediators of acquired resistance. Perhaps relatedly, effective therapies to circumvent or reverse acquired resistance largely remain elusive.

The landscape of immune acquired resistance to PD-(L)1 blockade is poorly understood. By contrast, several molecular mechanisms of acquired resistance to molecularly targeted therapies (e.g. EGFR and ALK-directed tyrosine kinase inhibitors) have been identified and led to significant therapeutic advances (Drilon et al., 2019; Piotrowska et al., 2018; Ramalingam et al., 2018; Shaw et al., 2019; Solomon et al., 2018). In patients with lung cancer treated with PD-(L)1 blockade, there have been a few published cases of acquired resistance (Abdallah et al., 2018; Anagnostou et al., 2017; Ascierto and McArthur, 2017; George et al., 2017; Gettinger et al., 2017; Iams et al., 2019; Koyama et al., 2016). Along with case reports in other diseases, these studies have identified that loss of key proteins associated with antigen presentation or defects of the IFNγ signaling pathway can contribute to immune resistance (Gettinger et al., 2017; Le et al., 2017; Sade-Feldman et al., 2017; Zaretsky et al., 2016). Pre-clinical work has further highlighted how the relative acuity vs chronicity of IFNγ exposure can contribute to immune dysfunction and tumor resistance (Benci et al., 2016, 2019; Grasso et al., 2020). Improved understanding of the nature and biology underlying acquired resistance is imperative to develop more effective next-generation immunotherapies in the future.

To address the clinical and molecular landscape of acquired resistance to PD-(L)1 blockade in patients with NSCLCs, we examined the largest clinical cohort (n = 1,201) of acquired resistance to PD-(L)1 blockade in lung cancer to date paired with a systematic genomic and transcriptomic analysis in a subset of patients (n = 29) with available tissue samples. We then also examined a murine model of acquired resistance to PD-(L)1 blockade to validate relationships identified in human samples.

## Results

### Acquired resistance to PD-1 blockade in NSCLC is common

Of 1,201 NSCLC patients treated with PD-1 blockade at MSK between April 2011 through December 2017, 243 (20%) achieved initial response. Many responding patients ultimately developed acquired resistance (AR), with an estimated cumulative AR rate of 61% (95% CI 36% - 85%) at 5 years of follow up using a competing risk model (**Figure 1a**). The onset of AR was variable (52% within 1 year, 39% 1-2 years, 11% >2 years) (**Figure 1b**). The relative risk of developing AR decreased with longer duration of initial response (**Figure 1c**).

**Figure 1.**
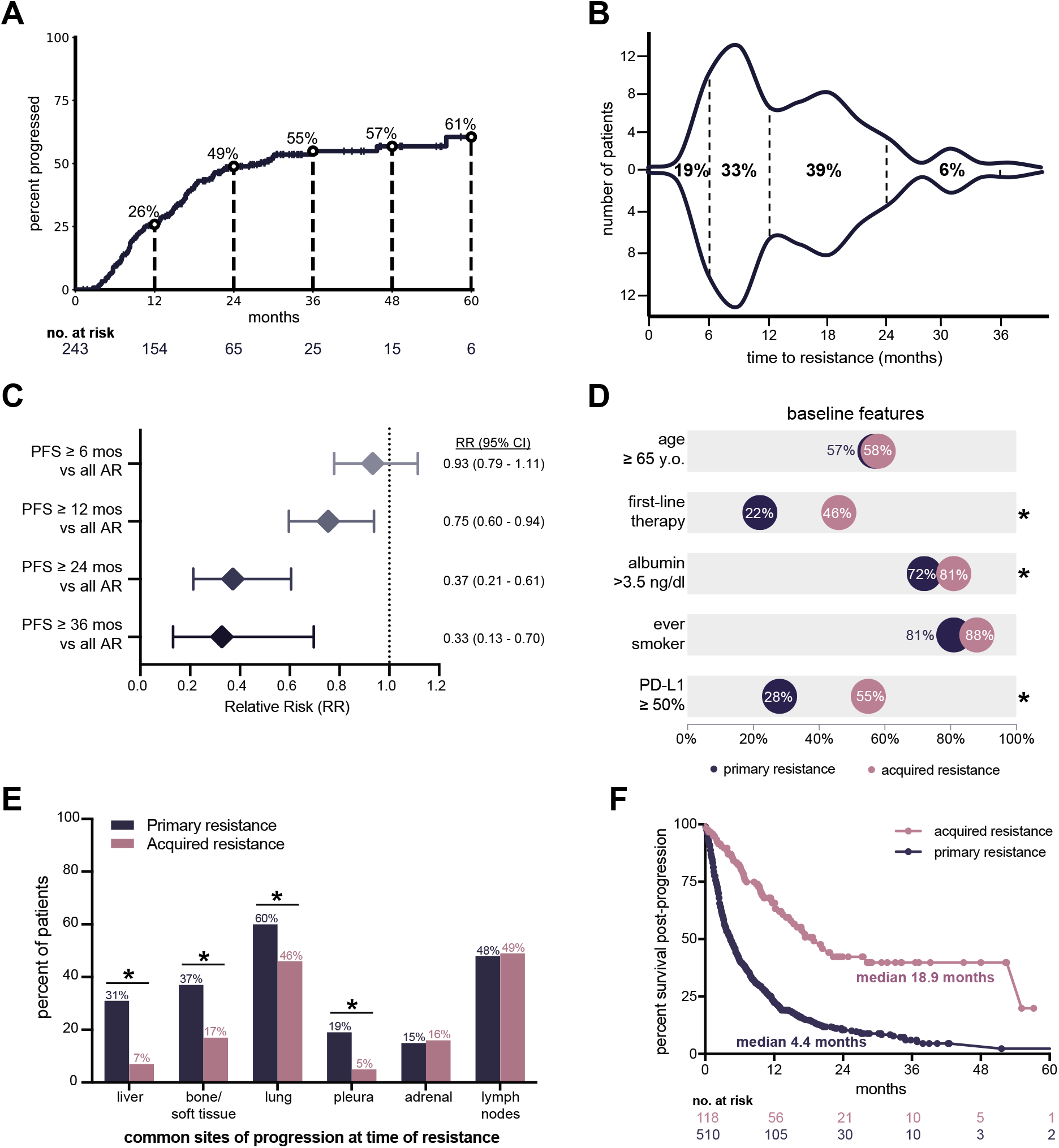
Clinical features of acquired resistance to immunotherapy in lung cancer. (a) Cumulative incidence of developing acquired resistance among NSCLC patients with initial response to PD-1 blockade therapy. (b) Time to onset of acquired resistance among responders. (c) Estimated rate of developing acquired resistance defined by duration of initial response. (d) Rates of baseline clinical features among patients with primary (n = 346) and acquired resistance (n = 118). Asterisk represents significant comparisons of p < 0.05. (e) Common organ sites of progression at time of primary or acquired resistance. (f) Post-progression overall survival in patients with primary or acquired resistance (p < 0.0001).

Although acquired and primary resistance have not been directly compared previously, we hypothesize that these scenarios are distinct biologically and clinically. Consistent with this, we found that several baseline clinical features differ between patients with acquired and primary resistance (**Figure 1d**). High tumor PD-L1 protein expression in baseline (pre-treatment) tissue, in particular, is enriched among patients with acquired resistance compared to primary resistance (55% vs 28%, Fisher’s p = 0.02). The organ-specific pattern of progression also differed, with liver metastasis being a common site of progression at primary resistance but relatively uncommon in acquired resistance (31% vs 7%, Odds Ratio 6.23, Fisher’s p<0.0001, **Figure 1e**). Perhaps most notably, the post-progression overall survival was significantly longer in patients with acquired resistance compared to primary progression (median 18.9 months vs 4.4 months, Log-rank p< 0.0001 **Figure 1f**), suggestive of persistent, partially effective anti-tumor immune responses that permits prolonged survival even after the initial onset of acquired resistance. Overall, AR is largely characterized by distinct clinical features, suggesting acquired resistance may have underlying immunobiologic features that are distinct from primary resistance and in need of dedicated analysis.

### Patient cohort for molecular profiling of acquired resistance to PD-1 blockade

To investigate the molecular mechanisms of acquired resistance to PD-1 blockade in patients with NSCLC, we generated microarray-based whole transcriptome expression data, and whole exome sequencing (WES) data from pre and/or post-treatment tumors in a subset of patients. Patients with analyzed samples had similar baseline characteristics to those in the larger clinical cohort (**Table S1**). After QC and sample prioritization, the primary analysis of the molecular data focused on 49 tumor samples (13 pre-treatment, 29 post-treatment) from 29 patients for expression data and 34 tumor samples (15 pre-treatment, 19 post-treatment) from 22 patients for exome data (**Figure 2a**, **Table S2**). 13 patients had expression data available from both pre- and post-treatment tissue; 12 patients had exome data available from both pre- and post-treatment tissue. All post-treatment samples were obtained following radiographic progression to PD-1 blockade (median time from progression to sample collection 3.7 weeks, IQR 1.8-10.4) and prior to initiation of new systemic therapy (**Figure 2b**).

**Figure 2.**
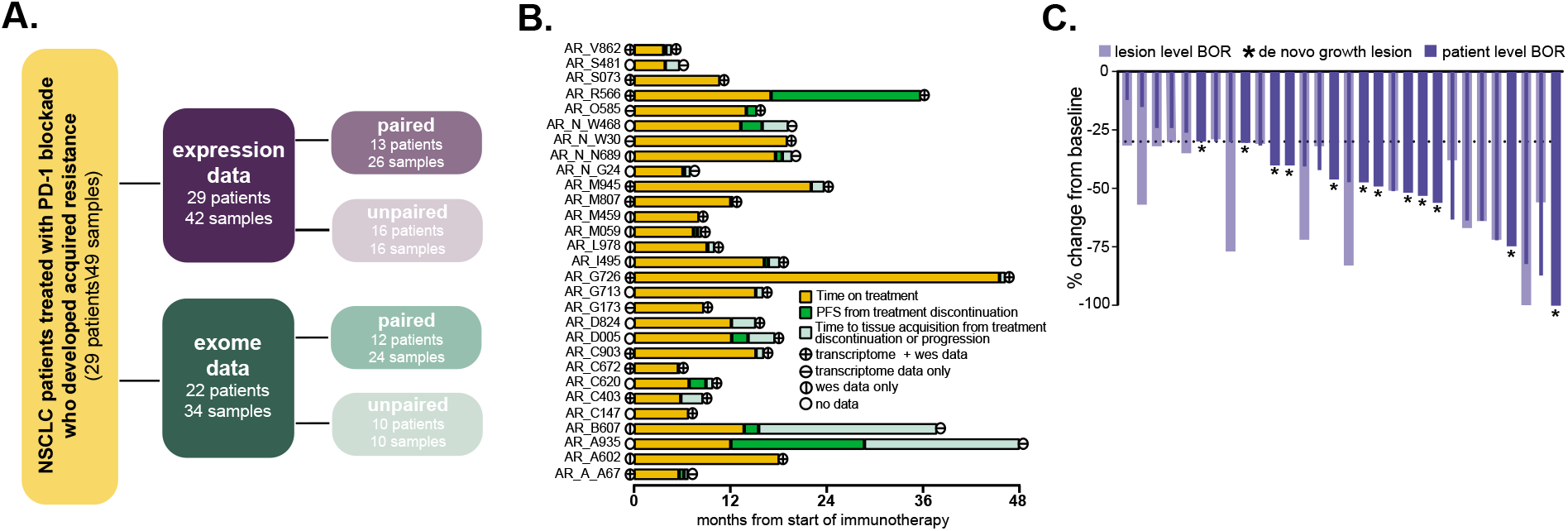
Overview of the patient cohort used for the exome and expression analyses. (a) Flow diagram depicting molecular profiling of samples from NSCLC patients treated with PD-1 blockade who developed acquired resistance. Paired samples are those collected prior to treatment initiation with PD-1 inhibitor and at time of resistance from the same patient. Unpaired samples include single timepoints of collection; prior to treatment initiation or at time of resistance. (b) Swimmer’s plot of when each patient was molecularly profiled. Course of treatment, progression-free survival, and time to tissue acquisition are depicted. Lines within circles identify the type of sequencing completed. (c) Waterfall plot of RECIST best overall response in patient (dark blue) and lesion (light blue). Dashed line represents 30% shrinkage. Asterisk represents new lesions that developed on treatment (de novo growth).

Our work and others (Gettinger, *J Thor Oncol* 2018) have shown that acquired resistance frequently occurs in an oligoprogressive pattern, highlighting the importance of assessing the lesion-level response in the analysis of acquired resistance. Therefore, we examined the lesion-level response (and resistance) from which each sample was collected to optimize that pre-treatment and post-treatment samples reliably represented the biology of responsive and acquired resistance tumors, respectively. Specifically, all post-treatment samples were derived from sites with lesion-specific radiologic rebound growth or de novo growth (**Figure 2c**, **Figure S1,S2**).

### Acquired resistance to PD-1 blockade is associated with a distinct transcriptional landscape

Principal components analysis (PCA) of protein-coding gene expression profiles from whole transcriptome data of all 42 samples showed no major technical or clinical factors influenced clustering, including batch and site of sample collection (i.e. lung, lymph node, adrenal, etc) (**Figure S3a,b**). There was also no separation among phenotypically distinct post-treatment lesions (**Figure S3c**). We summarized gene expression values to pathway-level scores using single-sample gene set enrichment approach (ssGSEA) (Hänzelmann et al., 2013) on hallmark gene sets categorized into oncogenic, cellular stress, immune, stromal and other processes as previously applied (Jiménez-Sánchez et al., 2020). PCA clustering of 24 paired samples using enrichment scores showed a separation of samples based on paired pre- and post-treatment timepoints, with the separation primarily driven by immune-related hallmark gene sets (**Figure 3a,b**). Differential expression analysis of paired samples for hallmark gene sets showed a significant upregulation of IFN alpha/gamma response, oxidative phosphorylation, and DNA repair pathways after treatment (FDR < 0.1, **Figure 3c**, **Table S3**). Clustering of paired samples based on computational deconvolution of immune cell estimates from bulk expression derived using CIBERSORT (Newman et al., 2015) showed a separation of pre- and post-treatment samples particularly driven by infiltration of CD8+ T cells (**Figure 3d,e**). Significant increase in immune infiltration (wilcoxon signed-rank test p<0.05; **Figure S3d**) and specifically CD8+ T cells was also observed post-therapy from differential analysis of paired pre-treatment and post-treatment samples (FDR < 0.1, **Figure 3f**, **Table S3**).

**Figure 3.**
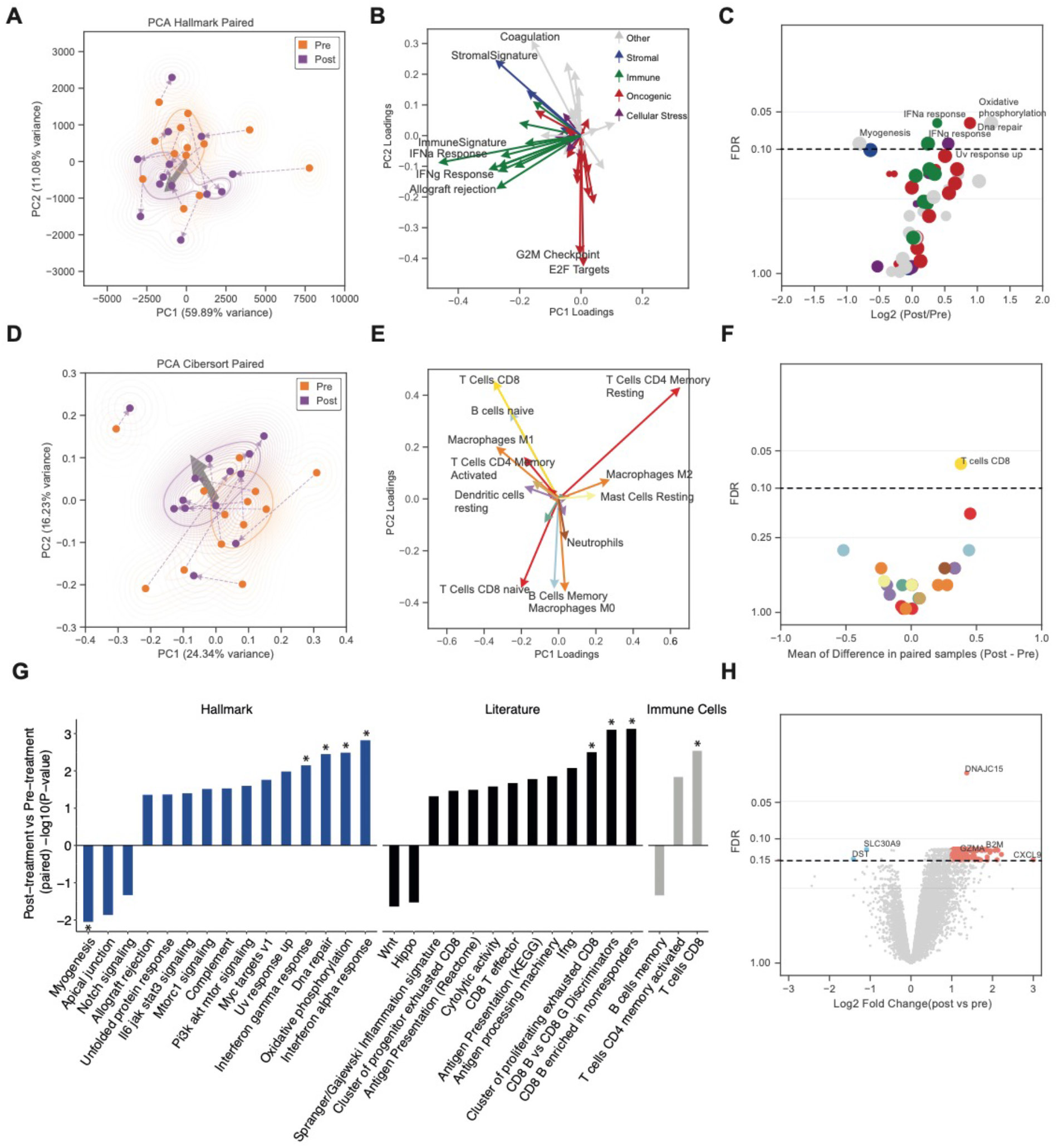
Resistant lesions show up-regulation of Interferon Gamma Response pathway and infiltration of CD8+ T cells. (a) Principal components analysis of paired samples using enrichment scores of hallmark gene sets derived from ssGSEA. Paired pre- and post-treatment lesions from the same patient are connected using a dashed line. The light grey arrow indicates the average directionality of change for each pair. (b) Principal components feature loadings of hallmark gene sets with both magnitude and direction. Biological processes in hallmark gene sets were categorized into sub-groups as described in (Jiménez-Sánchez et al., 2020) and colour-coded accordingly. (c) Differential comparison of hallmark enrichment scores (ES) between pre- and post-treatment samples. Each point represents a hallmark gene set and point size indicates the number of genes in a gene set. The x-axis indicates the change in hallmark enrichment scores for paired samples from each patient (Post vs Pre) and the y-axis is FDR adjusted p-value derived from the comparison of enrichment scores of hallmark gene sets using paired t-test. The black dashed line represents FDR cutoff to identify significant gene sets (FDR < 0.1). (d) Principal components analysis of immune cell estimates derived using CIBERSORT immune cell deconvolution approach. (e) Principal components feature loadings of immune cell estimates. (f) Differential comparison of immune cell estimates (CIBERSORT) between pre- vs post-treatment samples. Each point represents an immune cell type and associated colour reference indicated in panel e. The x-axis indicates the change in immune cell estimates for paired samples from each patient (Post vs Pre) and y-axis is FDR adjusted p-value derived from paired comparison of immune cell estimates. (g) Summary of key changes in hallmark gene sets, ICB-resistance related gene signatures and estimates of immune cells using differential analysis of expression data. All gene sets with p value < 0.05 are shown. * indicates gene sets that were significant after FDR correction (FDR < 0.1). (h) Differentially expressed genes between pre- and post-treatment samples. The black dashed line represents FDR cutoff to identify significant genes (FDR < 0.15). Benjamini–Hochberg (BH) method was used for false discovery rate (FDR) correction.

Several clinical and pre-clinical studies have generated bulk or single-cell RNAseq datasets to identify gene sets associated with ICB resistance and T cell dysfunction. We curated a non-redundant resource of these gene sets (See **Methods** for details) and compared differential changes among the paired samples (**Table S3, S4**). Among these, comparing post-treatment to pre-treatment samples, we found an increase in expression of antigen presentation (AP) pathway, IFNγ (Gao et al., 2016), CD8 T effectors (Rosenberg et al., 2016), and proliferating exhausted CD8+ T cells (Miller et al., 2019), while genes belonging to WNT (Sanchez-Vega et al., 2018) pathway showed modest reduction in expression (**Figure 3g**). Consistent with these gene sets associated with ongoing immune response to PD-1 blockade, expression of individual genes enriched in post-treatment tumors included *GZMA*, *B2M*, and *CXCL9* (**Figure 3h, Table S5**).

### Chronic and therapy-dependent increase in IFNγ response pathway as a potential route to acquired resistance to ICB

As sustained cancer-intrinsic IFN signalling has been linked to ICB resistance in pre-clinical mouse models of melanoma and other cancers (Benci et al., 2016, 2019; Jacquelot et al., 2019), we tested whether the change in ISG signatures (IFN*α* and IFNγ response) observed in our clinical cohort related to a resistance signature derived from a ICB-resistant mouse model of melanoma (Benci et al., 2019; Twyman-Saint Victor et al., 2015). We found a significant association between the mouse-derived ICB resistance signature and the treatment-induced change in IFNγ response (spearman’s rank correlation r = 0.90; p = 2.2e-16; **Figure 4a**), which persisted after removing overlapping genes (r = 0.86; p = 0.0003). Separately, principal components analysis of change in enrichment score of hallmark gene sets between paired lesions showed a separation of patients on the 1st principal components based on the extent of change of ISG signatures (**Figure S4a–c**). The correlation was significantly stronger for change in the IFNγ-specific response genes (r = 0.9; p < 2.2e-16) when compared to change in IFN±-specific response genes (r = 0.48; p = 0.09; **Figure S4d–e**). Notably, samples could be separated into two subsets, with about half of the paired samples showing no increase pre- to post-treatment and the other half characterized by induced expression in the IFNγ response pathway. This led us to categorise the samples into an IFNγ “stable” and an IFNγ “increase” group (**Figure 4b**).

**Figure 4.**
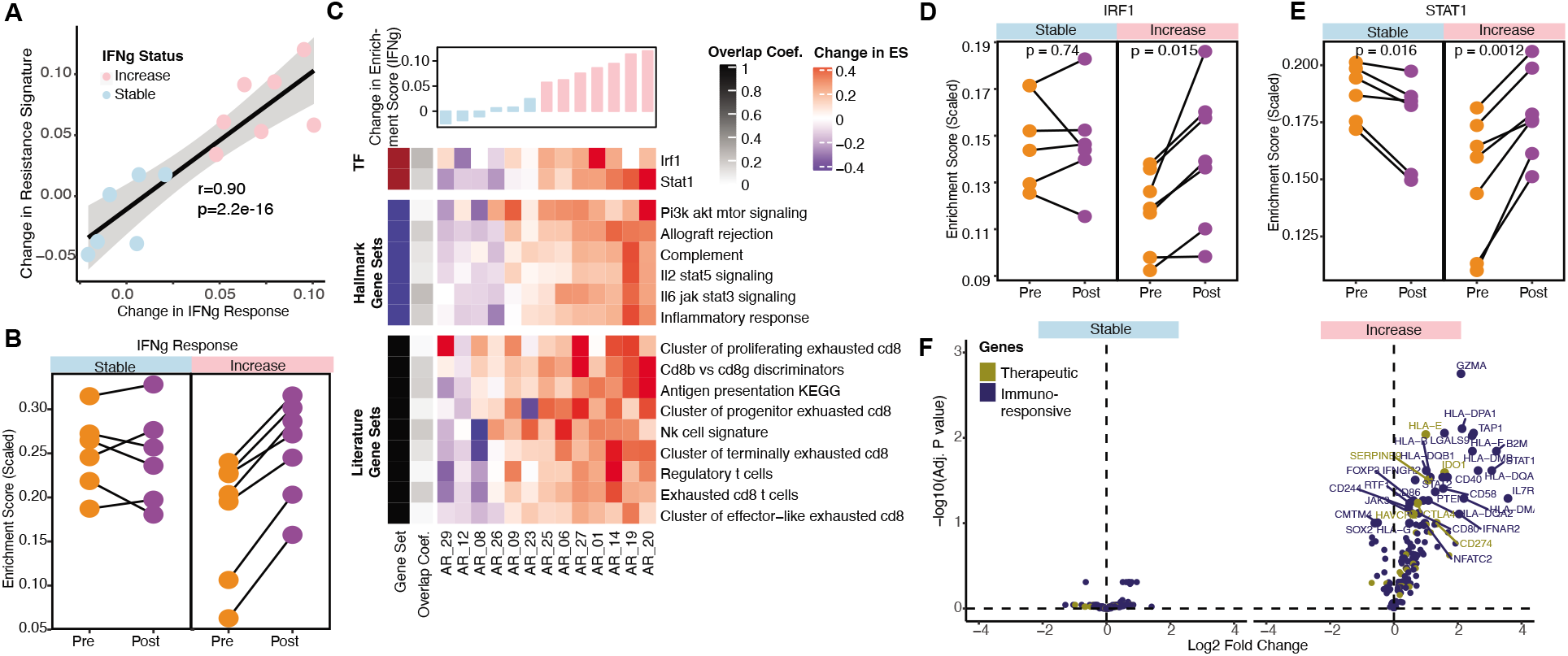
Activation of IFNγ correlates with CD8+ T cell exhaustion signature. (a) Correlation between change in the IFNγ response signature and change in the ICB-resistance signature derived from a mouse model of melanoma for the paired samples (Twyman-Saint Victor et al., 2015). (b) Patients were sub-divided into ‘stable’ and ‘increase’ categories based on the magnitude of change in the INFg response signature between the pre- and post-treatment samples. (c) Change in enrichment scores of key differentially regulated gene sets in either ‘stable’ or ‘increase’ patients (p < 0.05) ordered according to change in enrichment score of IFNγ response signature. Activity of IFNγ response associated transcription factors (d) IRF1 and (e) STAT1 in pre- and post-treatment timepoints of patients in ‘stable’ and ‘increase’ sub-groups. (f) Differential change in expression levels of previously reported immune-responsive genes and resistance associated therapeutic targets in literature in the ‘stable’ or ‘increase’ sub-group. Benjamini– Hochberg (BH) method was used for p-value adjustment. Statistical comparisons in panel d and e were performed using two-tailed paired t-test.

Consistent with a differential change of IFNγ in these patients, patients with increase in IFNγ generally had an increase in inferred activity of individual transcription factors associated with activation of IFN-stimulated genes (ISGs), *STAT1* and *IRF1* as well as immune signatures (estimated by hallmark and literature gene sets) associated with CD8+ T cell exhaustion across several studies (Miller et al., 2019; Sade-Feldman et al., 2018) (**Figure 4c–f, Table S6**). In addition to signatures of T cell exhaustion, increase in regulatory T cells (Sade-Feldman et al., 2018) were also noted. FOXP3 was specifically upregulated in the IFNγ ‘increase’ subgroup (paired t-test p = 0.005; **Figure S4f**). In contrast, patients with ‘stable’ IFNγ were characterized by a lack of change in these immune-related pathways and genes (**Table S6)**. Together these data suggest a recurrent pattern of acquired resistance to PD-1 blockade in NSCLC is characterized by activation of IFNγ transcriptional program in tumors, presumptive tumor-specific IFNγ insensitivity (given persistent tumor growth clinically), and a concomitant increase in exhaustion of CD8+ T cells in the micro-environment.

### Positive selection pressure for antigen presentation gene mutations in acquired resistance

To examine somatic alterations and potential mechanisms of acquired resistance, we next turned to evaluate the exome sequencing data pre vs post-treatment for 12 patients (24 samples with germline SNPs confirming paired samples belonged to the same patient; **Figure S5a**). NSCLC is characterized by a high mutation burden, a strong predictor of response to immunotherapy (Rizvi et al., 2015, 2020; Strickler et al., 2021). Overall, there was no significant difference in tumor mutation burden (wilcoxon signed-rank test p = 0.6; **Table S7**), known driver genes (Campbell et al., 2016), neoantigen burden, fitness (wilcoxon signed-rank test p = 0.74), or tumor heterogeneity (wilcoxon signed-rank test p = 0.37) before versus after immunotherapy treatment at a summary level (**Figure 5a**, **Figure S5b,c**). However, there was evidence of remodeling of clonal or sub-clonal structure in seven patients. For five of these patients, clonal mutations were retained while a subset of sub-clonal mutations were lost and/or new sub-clonal mutations were also acquired (**Figure 5b**, **Figure S5d**). For two patients (AR_20 and AR_27), post-treatment lesions did not share any somatic mutations with their respective pre-treatment lesions indicative of emergence of a potentially new tumour or outgrowth of a rare (i.e. below the limit of detection by WES) pre-existing tumor clone (**Figure S5e)**. Among the clonal mutations detected in post-treatment lesion of AR_20 included a nonsense mutation in STK11 gene consistent with previous observations of an association between mutations in STK11 gene and resistance to immune-checkpoint blockade in lung adenocarcinoma (Skoulidis et al., 2018). Several mutational processes including extrinsic factors, particularly smoking, can influence somatic molecular profile in NSCLC and can be detected as mutational signatures (Alexandrov et al., 2013). The smoking signature was the dominant signature in pre-treatment lesions and these mutations persisted in post-treatment lesions. However, post therapy the clonal composition of these tumours had changed potentially shaped by different sets of factors indicated by depleted proportion of smoking related mutations. (**Figure S5f**). Recent studies have shown an enrichment of APOBEC mutational signature in patients that benefit from immunotherapy treatment (Wang et al., 2018). In two patients, AR_08 and AR_20, we observed a noticeable increase in fraction of private mutations contributing to APOBEC mutational signatures 2 and 13 in the post-treatment lesions (48.3% in AR_08 and 14.3% in AR_20) relative to those in the pre-treatment lesions (5.4% in AR_08 and 1.6% in AR_20).

**Figure 5.**
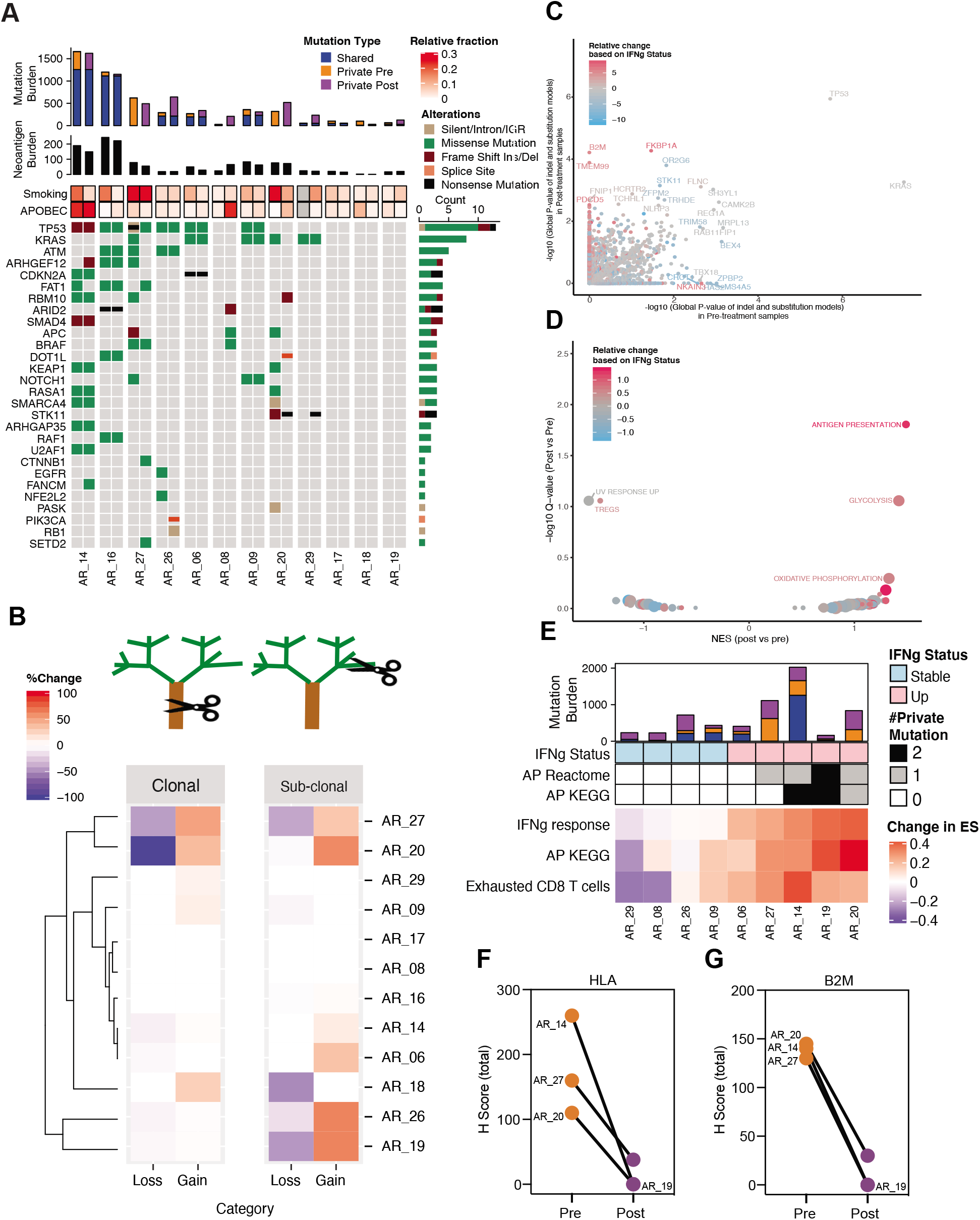
Genomic dynamics in acquired resistance to PD-1 blockade in lung cancer. (a) Summary of somatic mutations (missense and indels) in our ICB-resistance cohort for known driver genes in non-small cell lung cancer. Pattern of mutations of recurrently mutated genes derived from a previous study (Campbell et al., 2016). The heatmap also indicates the unique and shared mutations in each sample and the proportion of mutations associated with key somatic signatures (smoking and APOBEC) associated with lung cancer. (b) Percentage loss or gain of clonal and sub-clonal mutations in each patient. (c) Comparison of global p-value estimates for genes derived from dN/dS analysis of missense, truncations and indels to evaluate gene-level selection pressure in pre and post-treatment samples estimated using dndscv method (Martincorena et al., 2017). (d) Comparison of global p-value estimates genes to identify gene sets under positive selection in pre and post-treatment samples. The change in gene level global p-value between pre- and post-treatment samples (shown in c) was used to order genes and estimate GSEA normalized enrichment score and p-value for each gene-set. (e) Summary of key changes in expression and mutations in nine patients with pre- and post-treatment measurements for both expression and exome. The private mutations in post-treatment lesions of patients in genes part of Antigen Presentation Pathway (KEGG or REACTOME) are shown. Immunohistochemistry based quantification of (f) HLA/MHC-I and (g) B2M.

Given previous studies describing loss of B2M and other genes such as TAP1, TAP2 and TAPBP involved in antigen presentation pathway as a potential mechanism of immune escape in resistant tumours, we performed an unbiased analysis to evaluate positive selection pressure on individual genes before and after therapy (Martincorena et al., 2017). As expected, canonical driver mutations in lung cancer such as KRAS and TP53 were under strong positive selection pressure and there were no recurrently altered driver genes with significant enrichment in post-treatment tumors compared to pre-treatment (**Figure 5c, Figure S5g**). However, a nonsense mutation and a frameshift deletion in B2M were exclusively identified in post-treatment tumors of AR_14 and AR_19 respectively, and other immune-related genes such as IL21R, PDCD5, FKBP1A and FNIP1 were indeed enriched post-therapy (**Figure 5c, Figure S5h,i**). No potential pathogenic mutations were observed in TAP1, TAP2 and TAPBP genes.

Given the selective identification of mutation in B2M and other immune-related genes in the ICB-resistant tumor samples, we evaluated additional gene sets involved in Antigen Presentation (AP) pathways using the GSEA approach. Specifically, we asked whether there was evidence of an association between IFNγ selective pressure and dysregulation of AP pathways (**Figure 5d**). Overlaying mutational changes with IFNγ status for the cases with both expression and mutation data, we observed the mutation enrichment in the AP pathway to be more common among patients that show an ‘increase’ in IFNγ response in contrast to those with ‘stable’ IFNγ response pathway. Notably, 3 out of 4 patients with significant change in clonal or sub-clonal architecture (AR_20, AR_27, AR_19), also showed presence of new mutations in the AP pathway genes in their post-treatment lesions (**Figure 5e**). All 4 of these patients also had available tissue for B2M and class 1 HLA protein expression testing on tumor cells and all were negative or decreased from baseline (**Figure 5f,g**)

### Acquired tumor IFNγ insensitivity associates with ICB resistance

To further explore the transcriptional features that are associated with acquired resistance to ICB in our clinical cohort, we also examined cancer cell intrinsic transcriptional programs using a preclinical murine model system of acquired resistance to ICB inhibitors. Similar to PD-1-responsive human lung cancer, the CT26 murine model is carcinogen-induced, has high tumor mutation burden, and is highly sensitive to immunotherapy treatment (Zhong et al., 2020), and therefore well suited pre-clinical analogue for interrogating acquired resistance. As expected, after subcutaneous inoculation of CT26 cells, tumors showed significant reduction in tumor volume over 3 weeks of anti-PD-1 treatment (**Figure 6a**). To model acquired resistance, following anti-PD-1 treatment, persistent viable cells were excised, cultured *in vitro*, and reimplanted in mice. The re-implanted tumors were then re-treated and this process was repeated for several passages until the serially progressive CT26 tumors were no longer responsive to anti-PD1 antibody therapy (**Figure 6b**). Bulk RNAseq was performed on the ICB-resistant cancer cell lines derived from tumors from the 2nd round (n=2) and 4th round (n=4) of *in vivo* passage and compared against the ICB-sensitive parental cell line (n=3; **Figure 6c**). PCA of whole transcriptome data did not show any clear trend (**Figure S6a**), however, PCA of hallmark gene sets showed the parental and 2nd round samples tend to cluster separately from the 4th round samples with the separation mainly driven by Interferon Alpha/Gamma Response pathway (**Figure 6d,e**). Systematic comparison of 4th round samples with parental samples showed a significant upregulation of several biological processes including TNFalpha signaling and IFNα/IFNγ response pathway (FDR <= 0.1; **Table S8**). In contrast, no significant change in gene sets were observed from the comparison of the 2nd round and parental cell lines (**Figure S6b,c**). Alongside the increase in INFg response pathway in the 4th round cells, we also observed a correlated increase in STAT1 and IRF1 activity inferred from their regulons and expression of genes part of the Antigen Presentation Pathway (**Figure 6f–i**). Together, and similar to the human data, these results indicated that IFNγ response signaling was upregulated in ICB resistant cancer cells.

**Figure 6.**
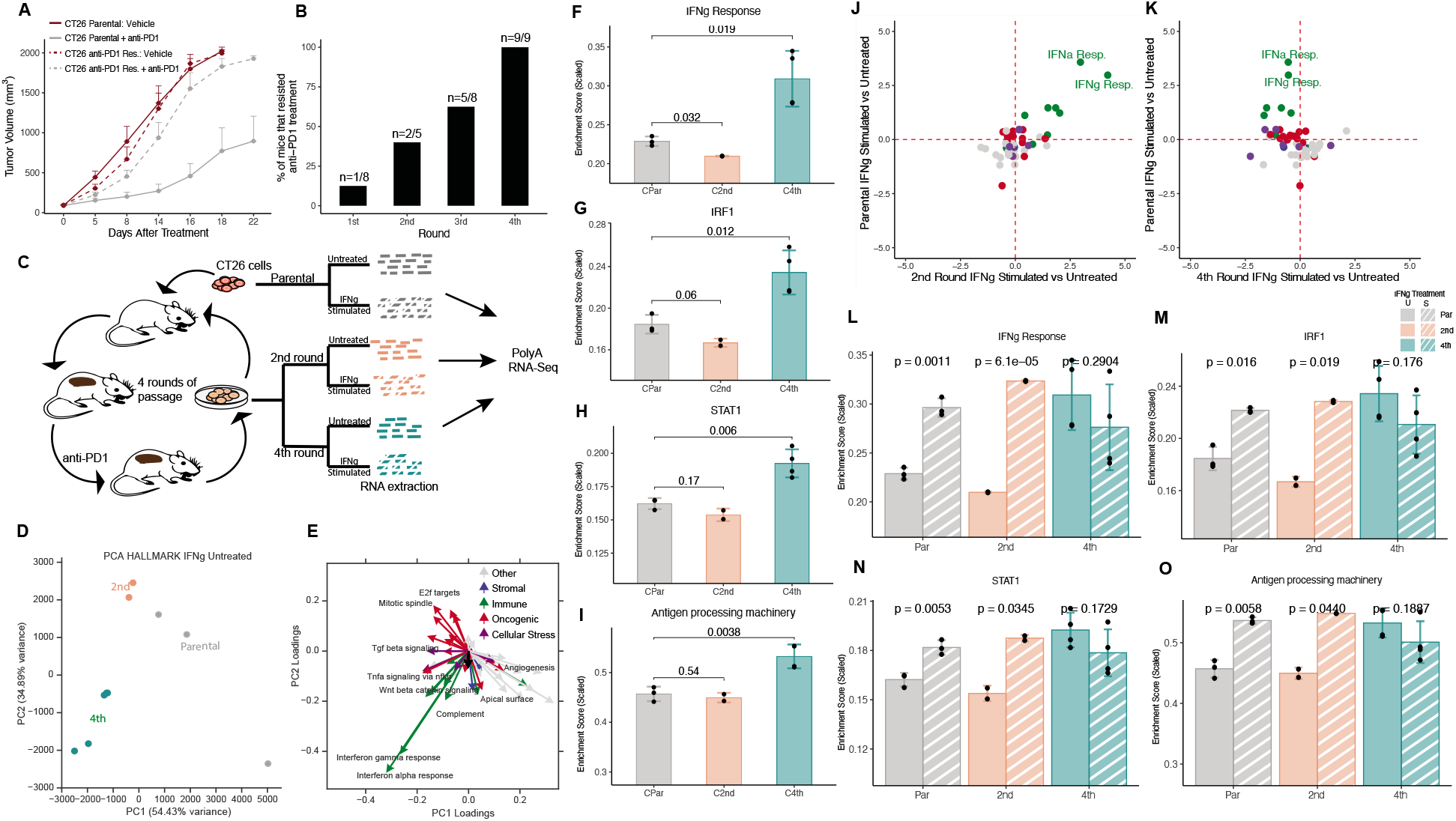
Cell lines derived from mouse CT26 tumours with acquired resistance to PD1 show dysfunctional IFNγ signalling. (a) Tumor volume over time after treatment with anti-PD1 therapy or control (Vehicle) for parental (CT26 parental) and resistant cells (CT26 anti-PD1 Res.) (n = 9 per group). (b) Percentage of mice that resisted anti-PD1 treatment. (c) Experimental design for development of ICB-resistance model from anti-PD1 treatment of CT26-derived tumours in mice. Cell lines were derived from tumors and subjected to RNA sequencing. (d) PCA of IFNγ-untreated samples i.e. parental (sensitive), 2nd round and 4th round ICB-resistant cells based on Enrichment Scores of hallmark gene sets. (e) Principal components feature loadings of hallmark gene sets with both magnitude and direction. Biological processes in hallmark gene sets were categorized into sub-groups as described in Jimenez-Sanchez et al, 2020 and the vectors were color-coded accordingly. Enrichment Scores in parental, 2nd and 4th round cells for the following genesets: (f) IFNγ response pathway, (g) STAT1, (h) IRF1 and (i) Antigen Processing machinery.(j) Comparison of significance of change in Enrichment Score between IFNγ Stimulated (IFNγs) and IFNγ Untreated (IFNγu) 2nd round and significance of change in Enrichment Score between IFNγs and IFNγu parental cells. (k) Comparison of significance of change in Enrichment Score between IFNγs and IFNγu 4th round and significance of change in Enrichment Score between IFNγs and IFNγu parental cells. Comparison of Enrichment Scores between IFNγu vs IFNγs (parental or 2nd or 4th) cells for the following genesets: (l) IFNγ response pathway, (m) STAT1, (n) IRF1 and (o) Antigen Processing machinery. Error bars are the standard error from the mean (n = 3 independent experiments). Statistical comparisons between parental and 2nd (or 4th) round samples or between IFNγs and IFNγu cells were made using two-tailed unpaired t-test.

To explore if ICB resistant cancer cells maintained responsiveness to type II IFN signaling, cell lines were stimulated with IFNγ for 24 hours and compared to unstimulated controls. While the parental and 2nd round cell lines showed an increased expression of genes involved in IFN signaling, the 4th round cell line did not show additional reactivity to IFNγ stimulation at the transcriptional level (**Figure 6j–l; S6d–f**; **Table S8**). In addition, downstream transcription factor activity of type II IFNγ signaling, such as STAT1 and IRF1, showed the same pattern with no statistically significant differences detected between IFNγ stimulated vs control in the 4th round cell line (**Figure 6m,n**). As type II IFN signaling is known to upregulate antigen presentation machinery pathway genes (Schroder et al., 2004), we also investigated the effect of IFNγ on these genes, which further supported the observations and showed no additional reactivity to IFNγ stimulation in 4th round cells (**Figure 6o**). Together these data indicate that 4th round PD-1 resistance cell lines have acquired insensitivity to IFN stimulation likely because they have high baseline IFN signaling activity and the pathway appears to be insensitive without canonical upregulation of interferon stimulated genes upon IFNγ exposure.

## Discussion

Although PD-1 blockade has been transformative in the treatment of patients with NSCLC, acquired resistance is common and understanding of the molecular mechanisms of resistance remains quite limited. Before embarking on this report, we had hypothesized that “non-inflamed” or “cold” tumors, characterized by exclusionary immunologic barriers or an absence of T cell infiltration, would significantly contribute to resistance (Schoenfeld and Hellmann et al., 2020). Previously, neoantigen loss and tumor-mediated immunosuppression have been associated with primary resistance to immunotherapy (Anagnostou et al., 2017; Peng et al., 2016; Verdegaal et al., 2016). In contrast, we found that neoantigen depletion does not appear to be dominant mediators of acquired resistance. In fact, most tumors have retained or increased inflammatory characteristics, rather than immune excluded or desert phenotype, with significant upregulation of IFNγ suggestive of persistent, albeit dysfunctional, anti-tumor immune response. The persistent, if incomplete, anti-tumor immune response may also manifest in the clinical observation that some patients who develop acquired resistance can still have durable survival for many years following initial emergence of resistance. In addition to the chronic upregulation of the IFNγ response pathway, we also observed strong upregulation of OxPhos and DNA repair pathway genes which is consistent with a recent report (Jaiswal et al., 2020) which proposes acquisition of hypermetabolic state with high expression of glycolytic and oxidative phosphorylation pathway genes as a potential escape mechanism in ICB-resistant melanoma cells.

The inflammatory phenotypes we identify have implications for future rational development of new immunotherapy strategies for patients with acquired resistance. Most notably, immune recruitment and infiltration did not appear to be the primary biologic challenges, which provides credence to strategies aimed to reprogram and rescue native anti-tumor immunity. Delivery of de novo anti-tumor immunity via engineered antigen-specific cellular or TCR-based therapies (Sarnaik et al., 2020; Creelan et al., 2020; D’Angelo et al., 2018; Doran et al., 2019; Nagarsheth et al 2021) also appears well-suited to exploit the lack of barriers to immune trafficking and persistent tumor antigen expression. While we did observe a few instances of sub-clonal/clonal neoantigen loss, these changes were relatively uncommon and mutation burden was generally unchanged pre vs post-treatment. One potential limitation of our clinical cohort is that it relies on bulk exome and transcriptome data which are prone to be affected by tumor purity. Future efforts utilizing single cell multi-omics will be important to further parse cancer cell-intrinsic vs immune or stromally-related mechanisms of resistance.

Our work informs and builds upon the prior pre-clinical and translational data supporting the intricate role of IFNγ in sensitivity and resistance to immunotherapy. Whereas initial IFNγ exposure may be fundamental to T cell activation and a hallmark of immune response, persistent IFNγ related effects and upregulation could signal immune dysfunction (Benci et al., 2016; Benci et al., 2019) and IFNγ insensitivity (Zaretsky et al 2016, Kalbasi et al. 2020, Grasso et al 2020). In contrast to previous reports linking IFNγ insensitivity to mutations in the JAK-STAT pathway, we did not identify specific defects in the IFNγ signalling pathway to explain the dysfunctional nature of IFNγ response observed in a subset of patients. While we found some evidence of B2M and other antigen presentation alterations, these changes were predominantly sub-clonal and generally co-occurred in tumors with upregulation of IFNγ potentially suggestive they are an evolutionary consequence rather than an initiating cause of resistance. We have previously shown that chronic IFNγ signaling may trigger a cascade of epigenetic modifications in tumor cells including enhanced IFN stimulated genes and ultimately generate a feedback loop of innate and adaptive immune exhaustion and dysfunction (Benci et al., 2016; Benci et al., 2019). In a murine model of acquired resistance presented here, we recapitulate how acquired resistance is associated with upregulated cancer-intrinsic IFNγ response and ultimately tumor insensitivity to effective anti-tumor immunity. Further work is needed to identify the specific mechanistic deficits in response to the dynamics of IFNγ signaling in both immune cells and tumor cells. Overall, these data can further guide more rationally guided therapeutic strategies to prevent, overcome, and reverse acquired resistance to PD-1 blockade for patients with lung cancer.

## Methods

### Description of the clinical cohort

Following MSKCC institutional review board approval, patients with advanced NSCLC treated with PD-(L)1 based therapy between April 2011 and December 2017 were identified. Response Evaluation Criteria in Solid Tumors (RECIST) version 1.1 was used to assess objective response outcomes. Patients with primary resistance were defined as those with progressive disease (PD) at their first on-treatment scan evaluation. Patients with acquired resistance were defined as those with partial or complete response (PR/CR) followed by isolated or systemic progression on or before the date of their last scan (median follow-up 33.6 months). Post-progression overall survival was calculated from the date of progression on PD-(L)1 inhibitor. Patients who did not die were censored at the date of last contact. A cumulative incidence function with death as a competing risk was used to estimate the proportion of acquired resistance over time. Overall survival was estimated using the Kaplan-Meier method.

### Generation of the molecular dataset

Tumor tissue samples from pre- and post-treatment timepoints were obtained from a subset of NSCLC patients treated with PD-1 blockade (n = 29). All samples were processed as FFPE. 16 samples were obtained prior to initiation of therapy (pre-treatment) and 37 samples were obtained at time of acquired resistance. Most patients had a best overall response of CR or PR per RECIST criteria (n = 22, 76%), with a small subgroup with stable disease (SD) less than −10%. Lesion-level response was obtained for all samples (**Figure S2**). Pre-treatment lesions were those that had at least a −30% reduction in size on treatment or were resected prior to initiating therapy (n = 4). In patients with resected lesion samples, lesion-level response could not be obtained so overall patient-level must have been CR or PR per RECIST. Of note, consistent with prior work demonstrating generalized inter-tumor uniformity of response, pre-treatment samples derived from resected tumors had similar molecular features of tumors in which the lesion-level response was known (**Figure S2**). All post-treatment samples were obtained following radiographic progression to PD-1 blockade. Post-treatment samples were defined as “rebound” or “de novo” growth (**Figure S1a**). Rebound lesions were those that were present at initiation of therapy, responded on treatment, but subsequently progressed. De novo growth lesions were those that were not present at initiation of therapy and newly grew following treatment. Time to progression in rebound lesions and growth of de novo lesions were similar (**Figure S1b**). Only patients with at least one post-treatment sample were included in this analysis. Samples were molecularly profiled by microarray-based transcriptome sequencing and/or whole exome sequencing. Expression and exome data were available on the same sample for 28 tumor lesions. The following antibodies were used for the Immunohistochemistry of the clinical : B2M (B2M Polyclonal, DAKO, A0072; RRID: AB_812325) and HLA (HLA-1/MHC-1, Clone: A4, eBioscience, 14-9958; RRID: AB_1210772).

### Processing and analysis of microarray data for the clinical cohort

Global RNA expression was measured using the human Affymetrix Clariom D Pico assay. The RNA samples quantification on Affymetrix Arrays was performed in two separate batches. Samples from each batch were processed independently using Affymetrix Expression Console Software. Initially the samples were normalized using the SST-RMA algorithm and outlier samples were excluded. Samples from the two batches were then combined together into a single dataset and subject to batch normalization using ComBat. Finally all the samples were further normalized together using LOESS normalization. For genes with multiple measurements, we selected the measurement with the highest coefficient of variation. The data analysis was focussed on 14,668 annotated protein-coding genes with expression measurements in the arrays. The expression dataset was thoroughly evaluated for technical artifacts such as batch effects (**Figure S3a**). Differential expression analysis was performed using the limma (Ritchie et al., 2015) package in R. Normalized expression data of protein-coding genes were fitted to a linear model using lmFit function and subject to empirical Bayes (eBayes) moderated t-statistics test to identify differentially expressed genes in paired lesions.

### Estimation of gene set enrichment scores from expression data

Enrichment scores were calculated for gene sets from the normalized expression matrix of protein-coding genes using the GSVA package (Hänzelmann et al., 2013) in R with default parameters except for method = ‘ssgsea’ and norm = ‘TRUE’. This approach was used to estimate enrichment scores for the Hallmark Gene sets (msigdb v6.1 database (Liberzon et al., 2015)) and non-redundant cancer and immune-related gene sets in literature. Although clinical data for acquired resistance to immunotherapy is fairly limited, recent studies have investigated the impact of chronic ICB treatment in *in vitro* cell lines and *in vivo* settings using mouse models generating either bulk or single-cell RNAseq datasets. We manually collated gene sets reported in many of these studies to build an extensive resource of biological processes and gene sets associated with cancer and immune pathways and more specifically ICB resistance (**Table S4**). In order to select for non-redundant gene sets from this resource, jaccard similarity coefficient was calculated between gene sets based on the number of shared genes and used this metric to perform hierarchical clustering of gene sets and construct a dendrogram with similar gene sets clustering together. Clusters of Gene sets were obtained by cutting the dendrogram at a particular level using cutree function in r (h=1.1). A non-redundant list of gene sets were created by selecting one gene set per cluster (**Table S4**).

Patients were classified into ‘increase’ and ‘stable’ sub-group based on the difference in the enrichment of IFNγ response pathway between the paired pre- and post-treatment samples. The ‘increase’ subgroup consisted of patients with difference in scaled enrichment score of IFNγ pathway > 0.025 while ‘stable’ subgroup was defined by minimal change in enrichment score of IFNγ pathway (< 0.025 and > −0.025). Overlap coefficients were calculated between IFNγ gene set and other gene sets to make sure correlation in enrichment score across samples was not driven by shared genes.Change in enrichment score in paired samples was calculated by first scaling the signed enrichment scores values using min-max normalization and then taking the difference in the scaled enrichment scores for paired post and pre-treatment samples for each patient. Enrichment scores were also calculated for transcription factors using the same approach as for other gene sets using previously published regulons of each of the 164 transcription factors (Garcia-Alonso et al., 2019). Deconvolution of immune cells from bulk microarray expression data was performed using CIBERSORT (Newman et al., 2015) tool with default parameters and normalized protein-coding gene expression matrix as input. Significance of change in enrichment score or immune cell estimates for paired samples was calculated using either paired t-test, welch t-test or wilcoxon signed-rank test depending on the evaluation of equality (Bartlett’s test or Levene’s test) of variance and normality assumptions. All pairwise correlations between gene sets based on change in enrichment scores were performed using spearman’s rank-order correlation method.

## Processing and analysis of Exome Data

### Processing of raw sequencing reads

Exome samples were aligned to human reference genome (hg19) using bwa aligner (v0.7.17) (Li and Durbin, 2009) and the aligned BAM files were subjected to deduplication and base recalibration methods in GATK (v4.0.2.1) (DePristo et al., 2011). These processed BAM files were used for all subsequent analyses.

### Detection of somatic mutations in tumor samples

Mutation calling for SNPs and Indels was performed for each tumor-normal(serum) pair using GATK-Mutect2 (v.4.0.2.1) with default parameters and additional filters to remove germline mutations including SNPs detected in gnomAD (Genome Aggregation Database) (Karczewski et al., 2020) and mutations in PoN (panel of normal) samples obtained from combining all normal samples in the cohort. Since exome samples were generated in multiple batches with different sequencers and using different capture kits (Illumina’s Rapid Capture Exome Kit (38Mb target territory), Agilent SureSelect Human All Exon V2 (44Mb target territory), Agilent SureSelect Human All exon V4 (51MB target territory)), mutations were only called on common regions captured by the three different kits. The mutation calls were annotated using the Oncotator v1 (Ramos et al., 2015) tool.

### Tumor heterogeneity and clonality

Tumor heterogeneity was evaluated using the Mutant-Allele Tumor Heterogeneity (MATH) score derived from the variant allele frequencies of somatic mutations as described previously (Mroz and Rocco, 2013). The clonal population structure of somatic mutations in tumor samples was inferred using Pyclone-VI (Roth et al., 2014). This method uses a bayesian statistical approach to estimate cellular prevalence of mutations after accounting for purity of samples. The tumor purity estimates for Pyclone-VI were obtained from FACETS (Shen and Seshan, 2016) and manually corrected for each sample based on the distribution of variant allele frequency. The mean cellular prevalence (MCP) estimates from Pyclone-VI were used to classify somatic mutation as clonal (MCP > 0.6), sub-clonal (MCP <= 0.6) or absent (MCP < 0.02).

### Estimation of somatic signatures in tumor exome data

Mutational signatures were estimated using the Sigfit (Gori and Baez-Ortega, 2018) package in R. For each exome sample, the proportion of mutations associated with each of the 30 mutational signatures in the COSMIC (Tate et al., 2019) database were estimated. Signature 4 corresponds to smoking signature while Signature 13 corresponds to APOBEC signature.

### Analysis of selection pressure in mutation data

Gene-level selection pressure was quantified for pre-treatment and post-treatment samples using the dNdScv (Martincorena et al., 2017) package in R. The dNdScv approach quantifies dN/dS ratios based on missense, truncations (nonsense and essential splice site) and indel mutations in a group of samples and identifies genes under positive selection in cancer based on the global p-values derived from likelihood tests. Selection pressure was calculated for each gene for pre-treatment and post-treatment samples separately and the difference in selection pressure between the two groups was used to identify potential biologically important genes associated with acquired resistance to ICB treatment. Gene sets with significant change in selection pressure between pre and post-treatment samples were identified via the GSEA (Subramanian et al., 2005) approach using the clusterProfiler (Yu et al., 2012) package in R with the difference in the - log10(global p-values) between pre and post-treatment samples, used as a metric to rank genes.

### Phylogeny Tree Reconstruction

For 12 patients with both pre and post-treatment WES available, mutations were filtered based on the following criteria: 1) total coverage for tumor ≥10, 2) variant allele frequency (VAF) for tumor ≥4%, 3) number of reads with alternative allele ≥9 for tumor, 4) total coverage for normal ≥7, and 5) VAF for normal ≤1% at a given mutation. These filters applied to all mutations except for mutations in the *KRAS* gene. Then pre- and post-therapy mutations were aggregated per patient. PhyloWGS (Deshwar et al., 2015) software package (https://github.com/morrislab/phylowgs) was used to infer the clonal structures and estimate clone sizes.

### Neoantigen Prediction and Fitness Score

Filtered mutations were annotated with snpEff.v4.3t software (Cingolani et al, 2012, PMID: 22728672) with options set as “ -noStats -strict -hgvs1LetterAa -hgvs -canon -fastaProt [fasta file name]”. All wild-type (WT) and mutant genomic sequences corresponding to coding mutations were translated to an amino acid sequence consistent with the GRCh37 reference genome (GRCh37.75). Only annotations without “WARNING” or “ERROR” were kept and the most deleterious missense mutation was prioritized in mapping a genomic mutation to a gene.

The mutant amino acid from a missense mutation was centered in a 17 amino acids long peptide. Then 9-mers were extracted in a left-to-right sliding fashion. Each mutant 9-mer contained the mutant amino acid on one of the 9 positions. In essence, one missense mutation produced up to nine 9-mer peptides. Predictions of MHC Class-I binding for both wildtype peptide (*P^WT^*) and mutant peptide (*P^Neo^*) were estimated using the NetMHC 3.4 (Lundegaard et al., 2008) software with patient-specific HLA-I types. All *P^Neo^*s with predicted IC50 affinities below 500 nM to a patient-specific HLA-I type were defined as neoantigens. Filtered neoantigens were aligned to the known positive epitopes in the Immune Epitope Database (IEDB, http://www.iedb.org) (Vita et al., 2019) for all human infectious disease, class-I restricted targets with positive immune assays using blastp (Altschul et al., 1990) software (https://blast.ncbi.nlm.nih.gov/Blast.cgi). We then calculate the alignment scores with the Biopython Bio.pairwise2 package (http://biopython.org) for all identified alignments.

Clonal structure, MHC Class-I affinities, and epitope alignment scores were put together into the fitness modeling framework in Luksza et al (Łuksza et al., 2017). Neoantigens were mapped to the clonal structure based on the underlying genomic mutations. Then fitness score was calculated for each clone, and the scores were averaged over all the clones in a sample after weighting on clonal sizes.

### Generation of anti-PD1 resistant CT26 tumors

BALB/C mice were acquired from The Jackson Laboratory, after several days of acclimation, mice were inoculated with 500k CT26 on the rear flank. When the average tumor volume reached 80-100 mm^3^ (indicating day 0), mice were given a series of intraperitoneal injections of anti-PD1 (clone RMP1-14; BioXcell), consisting of 100 ug each on days 0,3, and 6. Tumors were excised from mice that did not respond to anti-PD1 therapy, approximately 10-14 days following the initial treatment. Tumors were dissociated using collagenase (Stemcell Technologies), washed in 1X PBS, and plated in IMDM culture media supplemented in 10% fetal bovine serum, 1% GLUTiMAX, and 1% Antibiotic-Antimycotic (all GIBCO). Cells were passaged at least 5 times and then inoculated into new recipient mice according to the same protocol as above. Again, when tumors reached 80-100 mm3, another treatment course of anti-PD1 began. This process was repeated for a total of 4 rounds, at which point none of the treated mice responded to anti-PD1 therapy. The cell lines generated after 2 rounds of anti-PD1 selection are referred to throughout this manuscript as ‘2nd round’, ‘2nd generation’ or ‘F2 generation’, and the cell line generated after 4 rounds of anti-PD1 selection are referred to as ‘4th round’, ‘4th generation’, or ‘F4 generation’.

### Transcriptomic profiling of anti-PD1 resistant cell lines

Three distinct vials of parental CT26 cells (ATCC; ‘experimental replicates’), two independently isolated tumors from ‘2nd round’ mice, and four independently isolated tumors from ‘4th round’ mice (both ‘biological replicates’), were cultured +/−20 ng/ml of mouse IFNγ (Biolegend) for 24 hours at 37°C/5%CO2. The following day, RNA was isolated from cells using Qaigen RNAeasy reagents according to manufacturer’s instructions, including QiaShredder homogenization and on-column DNaseI digestion. Isolated RNA was sent to Genewiz (www.genewiz.com) for library generation, RNA-sequencing, and data processing. Briefly, sequencing libraries were generated and sequenced on an Illumina HiSeq (2×150 paired end reads), targeting >20×10^6^ reads per sample. Sequences were trimmed using Trimmomatic v.0.36 (Bolger et al., 2014) and mapped to Mus Musculus GRCm38 reference genome using STAR aligner v.2.5.2b (Dobin et al., 2013). Unique gene hits were calculated by using featureCounts from the Subread package v.1.5.2. Only unique reads that fell in exonic regions were counted. The Transcript Per Million (TPM) values were obtained for each protein-coding gene and subsequently log-transformed (log2(TPM +1)) for downstream analysis. Mouse orthologs of genes in hallmark gene sets and Antigen Processing Machinery (Şenbabaoğlu et al., 2016) and regulons of IRF1 and STAT1 were identified using Ensembl 87 (Howe et al., 2021) and single sample gene set enrichment analysis was performed in a similar fashion as described for the clinical cohort.

### Statistical tests of experimental data

All statistical tests were performed in R. The two-tailed unpaired t test was performed to identify differences between parental and the resistant cells or between naive and IFN*γ* challenged cells. Data are shown as mean + SEM.

## Supplementary Figures

**Figure S1.**
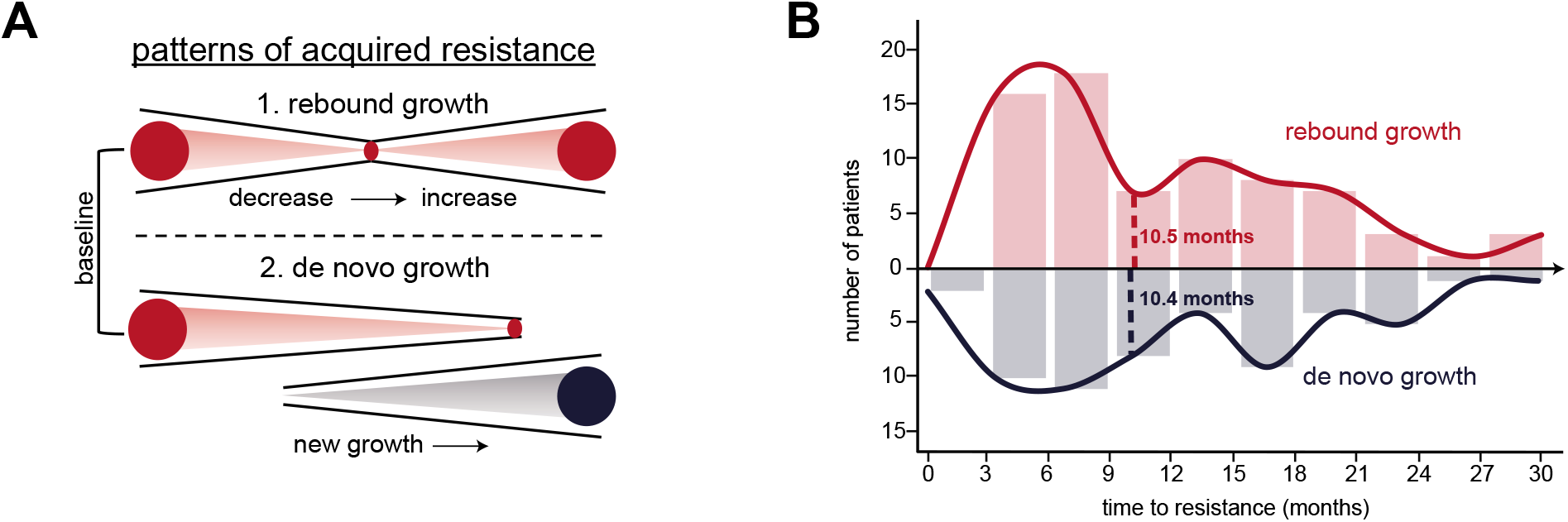
(a) Schematic of patterns of progression in post-treatment lesions. “Rebound growth” is defined as lesions present at initiation of therapy with response on treatment, but subsequently followed by progression (top). “De novo growth” is defined as lesions that were not present at initiation of therapy and newly grew during treatment. (b) Time to progression in rebound and de novo growth lesions.

**Figure S2.**
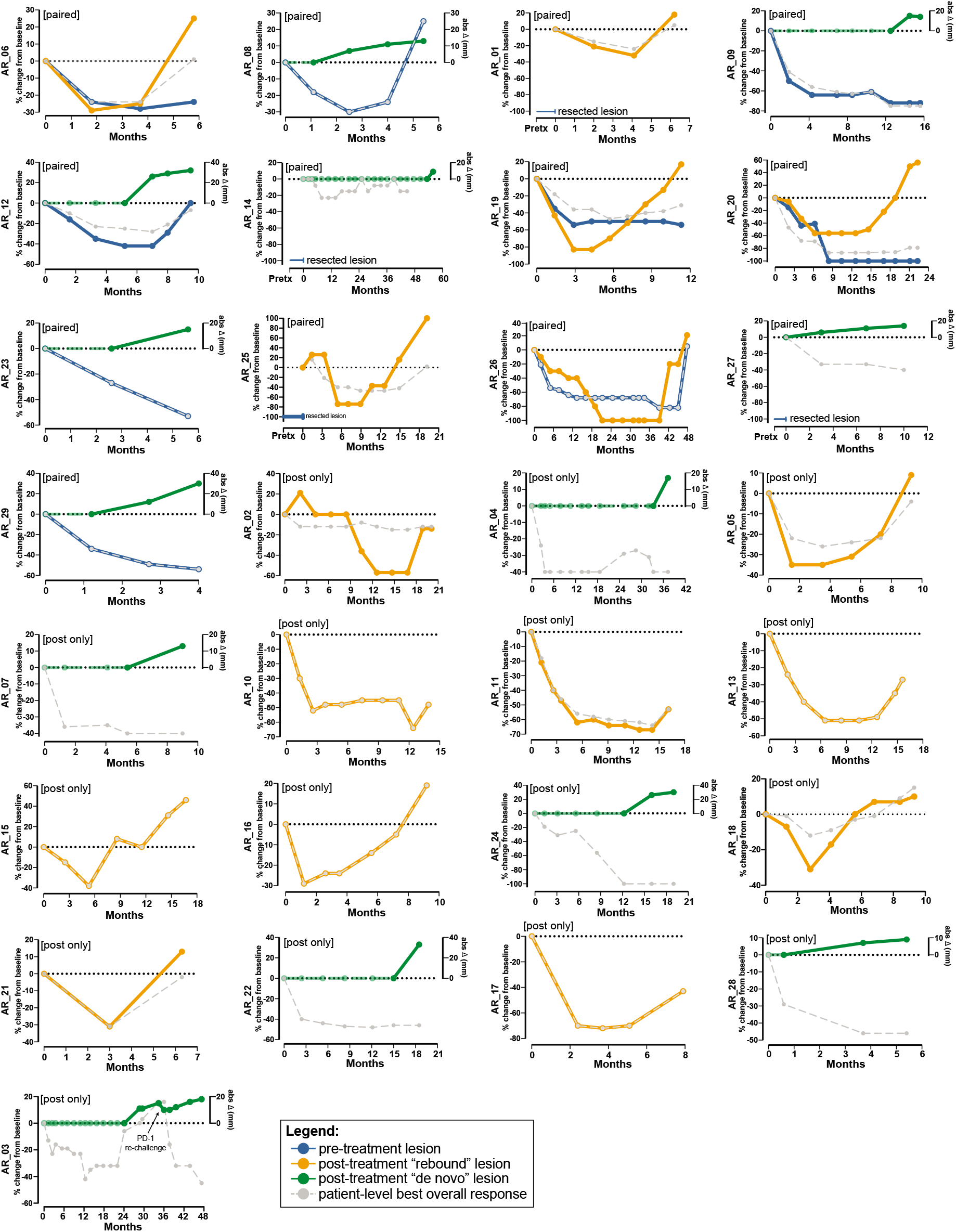
Spider plot of individual lesion and best overall response within each profiled patient. Pre-treatment lesions were defined as with at least 30% reduction after initiation of treatment or resected prior to treatment. Post-treatment lesions were defined as those with “rebound growth” or “de novo” growth.

**Figure S3.**
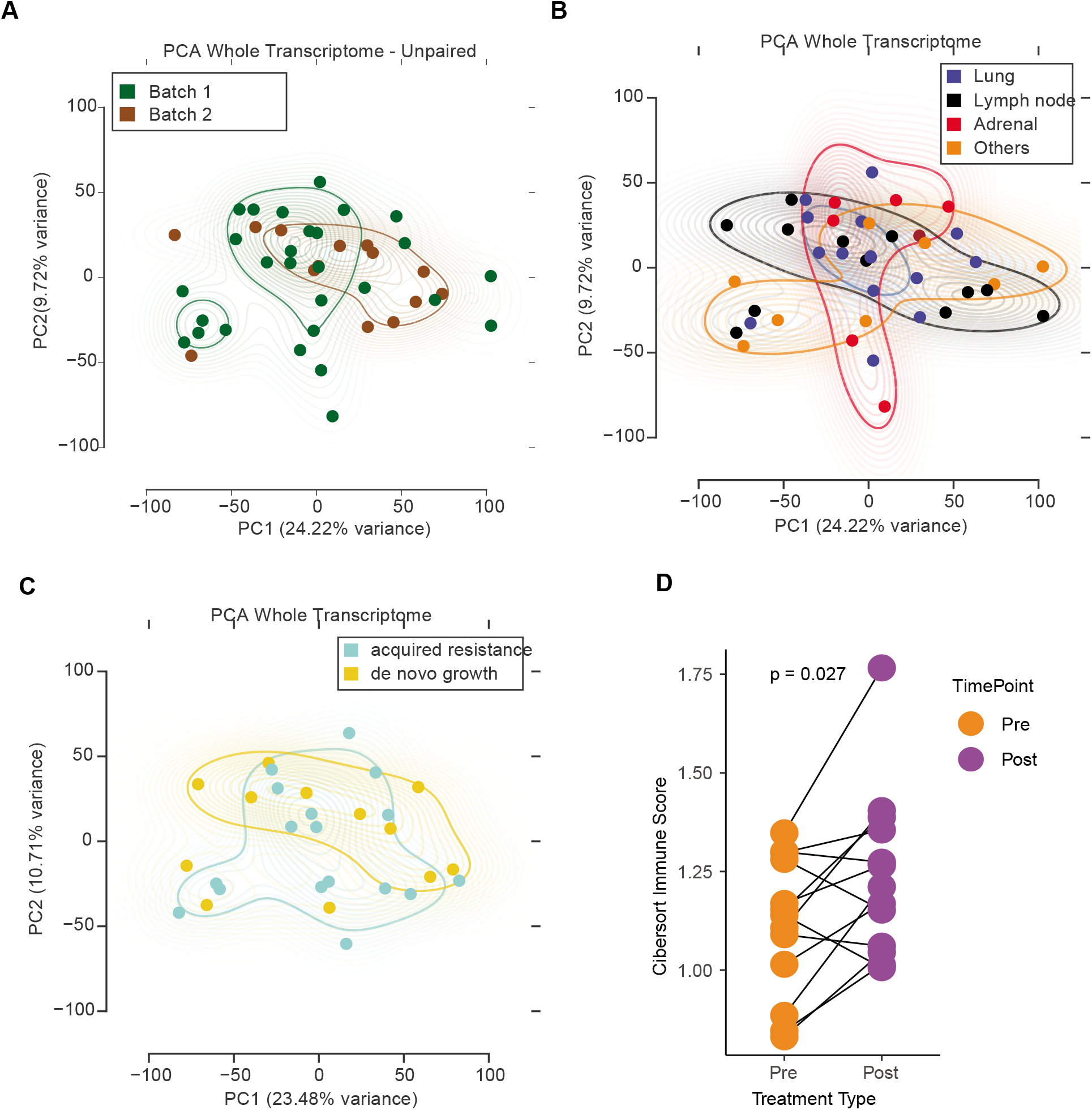
Principal components analysis of expression data to evaluate for a) batch effects, b) site and c) post-treatment response. d) Comparison of immune score from CIBERSORT between paired pre-treatment and post-treatment samples.

**Figure S4.**
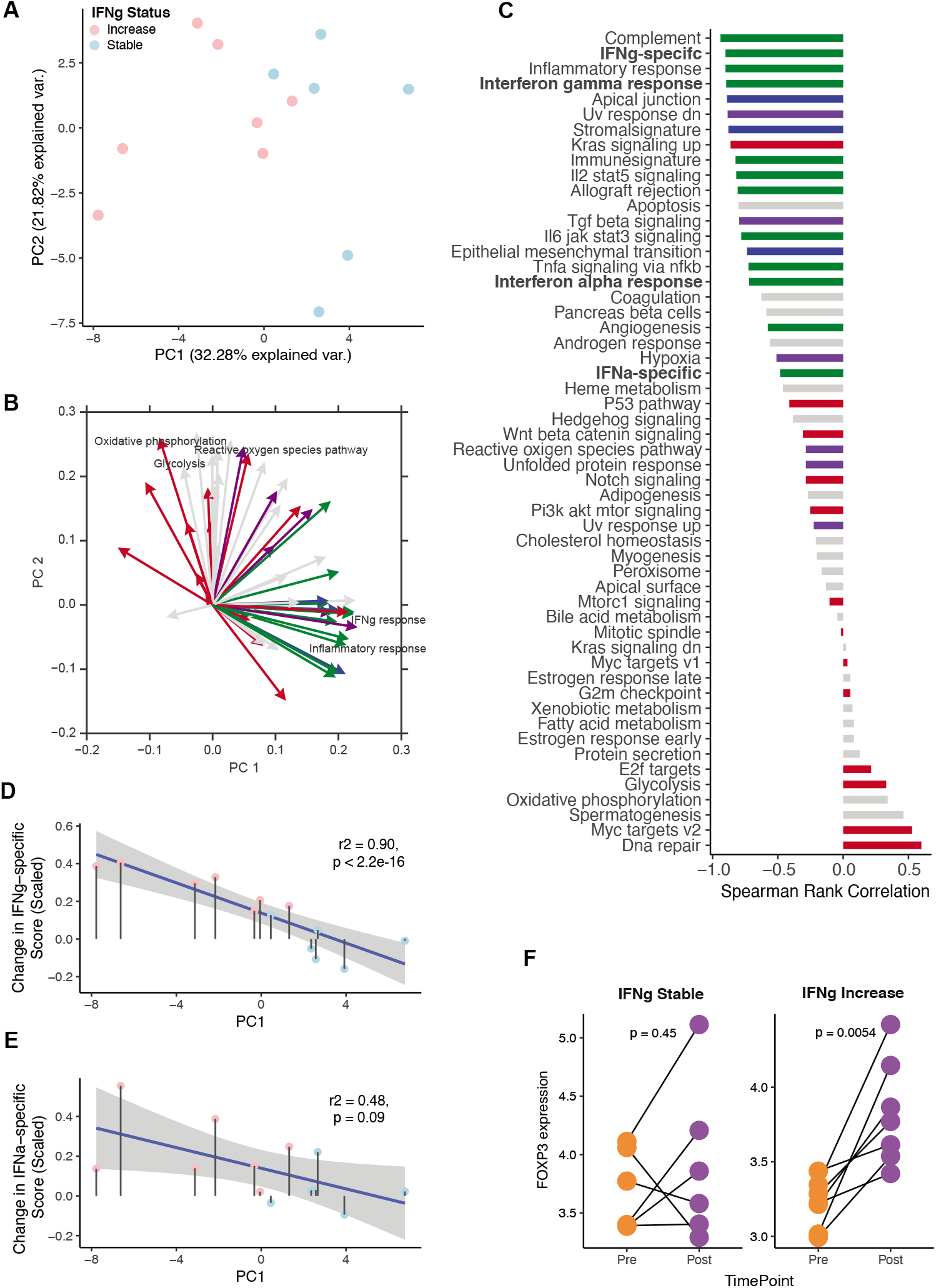
(a) Principal components analysis of patients with paired samples based on change in enrichment score of hallmark gene sets. (b) Principal components feature loadings of hallmark gene sets with both magnitude and direction. Biological processes in hallmark gene sets were categorized into sub-groups as described in (Jiménez-Sánchez et al., 2020) and colour-coded accordingly. regulated gene sets in either ‘stable’ or ‘increase’ patients ordered according to IFNγ status. (c) Correlation between Hallmark gene sets and PC1 from panel a.(d) Correlation between change in the IFNγ Response specific signature with the 1st principal components from panel a. (e) Correlation between change in the IFNα Response specific signature with the 1st principal components from panel a. (f) Expression levels of FOXP3 (Regulatory T cells marker) in pre- and post-treatment samples of IFNγ stable and increase group of patients.

**Figure S5.**
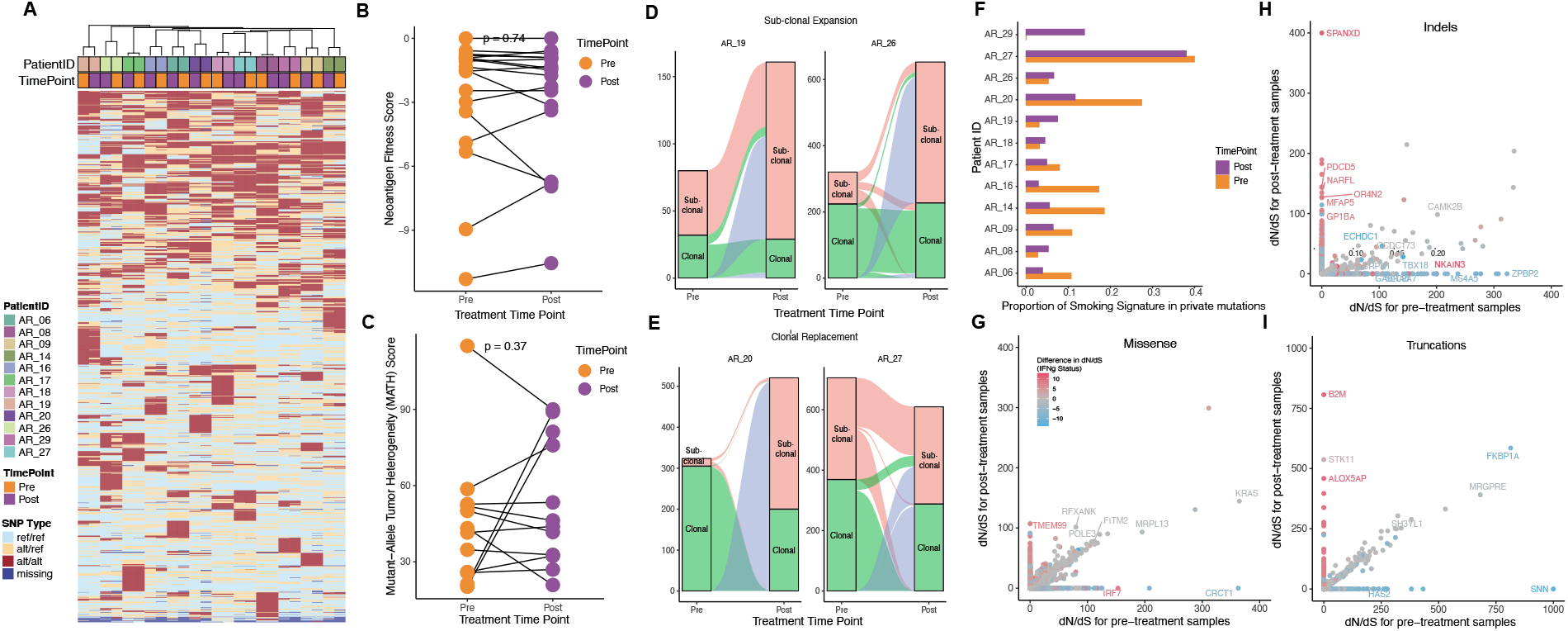
(a) Clustering of samples based on germline mutations. Column annotations indicate patient ID and treatment timepoints. Non-homozygous germline mutations were identified by genotyping the SNPs in the panel of normal (PoN) samples in the cohort using GATK. The variant allele frequencies of these germline SNPs in the tumour samples were used to classify them as ref/ref (<0.25), alt/ref (>=0.25 & <0.75) or alt/alt (>=0.75). (b) Neoantigen Fitness Score and (c) Mutant-Allele Tumor Heterogeneity (MATH) Score in paired pre-treatment and post-treatment samples. Dynamics of loss and gain of clonal and sub-clonal mutations between paired pre-treatment and post-treatment samples of patients with (d) sub-clonal expansion (AR_19 and AR_26) or (e) clonal replacement (AR_20 and AR_27). (f) Proportion of smoking signature in private pre-treatment and private post-treatment mutations. dN/dS values for genes based on (g) missense, (h) indel and (i) truncation (nonsense and splice site) mutations. Genes with high magnitude of dN/dS values (>100) and significant p-value (<.002) in either the pre-treatment or post-treatment timepoints are shown.

**Figure S6.**
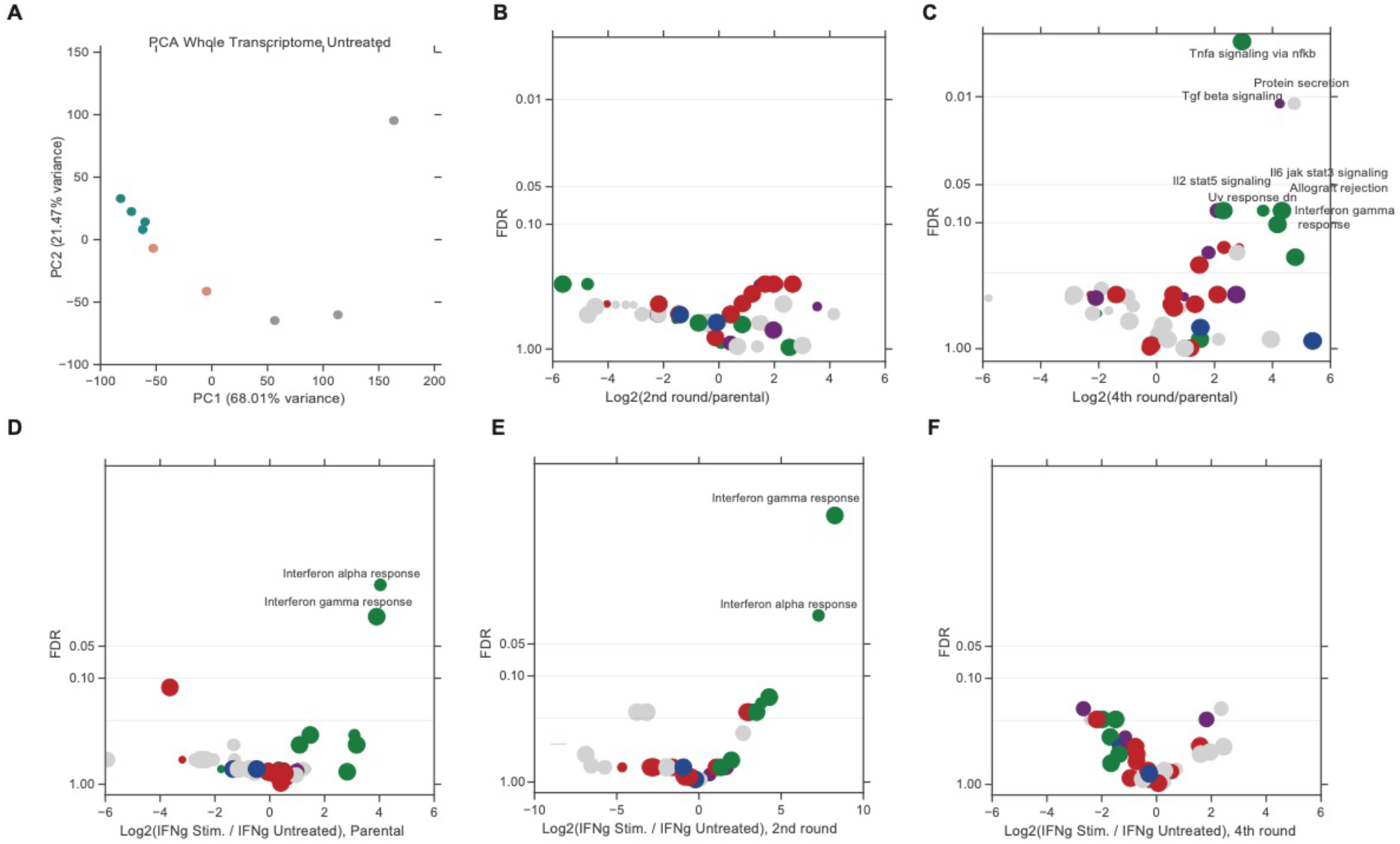
(a) PCA of parental, 2nd round and 4th round samples based on whole transcriptome data. Differential comparison of hallmark gene sets between (b) 2nd round and parental cells, (c) 4th round and parental cells. The x-axis indicates the change in Enrichment Score and y-axis is FDR adjusted p-value derived from comparison of Enrichment Score of hallmark gene sets. Differential analysis of hallmark gene sets for IFNγ Stimulated vs IFNγ Untreated cells from (g) Parental, (h) 2nd round and (i) 4th round.

